# The Urinary Microbiome in Postmenopausal Women with Recurrent Urinary Tract Infections

**DOI:** 10.1101/2020.12.21.423901

**Authors:** Monique H. Vaughan, Jialiang Mao, Lisa A. Karstens, Li Ma, Cindy L. Amundsen, Kenneth E. Schmader, Nazema Y. Siddiqui

## Abstract

Recurrent urinary tract infections (UTI) are highly prevalent in postmenopausal women, where vaginal estrogen and prophylactic antibiotics are used for treatment. The etiology of recurrent UTIs is not completely known, but the urinary microbiome is thought to be implicated. Thus, we aimed to compare the “steady state” urinary microbiome in three groups of menopausal women who were all using topically-applied vaginal estrogen: 1) women with recurrent UTIs *on* daily antibiotic prophylaxis; 2) women with recurrent UTIs *not on* antibiotic prophylaxis; and 3) age-matched controls without recurrent UTIs. Here we present a cross-sectional analysis of baseline data from 64 women enrolled in a longitudinal cohort study. Catheterized urine samples were collected > 4 weeks after last treatment for UTI. Samples were evaluated using expanded quantitative urine culture (EQUC) and 16S rRNA gene sequencing. With EQUC techniques, there were no significant differences in the median numbers of microbial species isolated among groups (p=0.96), even when considering *Lactobacilli* (p=0.72). However, there were trends towards different *Lactobacillus* species between groups. With sequencing the overwhelming majority of urinary samples contained *Lactobacilli*, with non-significant trends in relative abundance of *Lactobacilli* among groups. Using a Bayesian regression analysis for compositional data, we identified significant differences in anaerobic taxa that were associated with phenotypic groups. Most of these differences centered on Bacteroidales and the family *Prevotellaceae*, though differences were also noted in Actinobacteria and certain genera of Clostridiales. Associations between anaerobes within the urinary microbiome and recurrent UTI warrants further investigation.

**IMPORTANCE:** In menopausal women with recurrent urinary tract infections (UTIs) compared to those without, the abundance of *Lactobacillus* within the urinary microbiome is not significantly different when vaginal estrogen is regularly used. In this population, *Lactobacillaceae* were identified in 97% of urine samples using culture-independent techniques. However, with expanded urine cultures, women with recurrent UTIs taking daily antibiotics had a disproportionately low amount of *L. gasseri*/*L. acidophilus* compared to the other phenotypic groups. These findings support the theory that certain *Lactobacillus* species may be more important than others in the pathophysiology of postmenopausal recurrent UTIs. Furthermore, when using culture-independent techniques to explore urinary microbiota across phenotypic groups, we identified differences in multiple anaerobic taxa. Taken together, these results suggest that altered ratios of anaerobes and certain *Lactobacillus* species within the urinary microbiome may be implicated in postmenopausal recurrent UTI.

## INTRODUCTION

Recurrent urinary tract infections (UTIs), most commonly defined as 2 or more UTIs within a 6-month period, or 3 or more UTIs within one year (1), are common among women (2). Recurrent UTIs are particularly common in post-menopausal women (3), where the high prevalence of asymptomatic bacteriuria (4) and lack of diagnostic precision makes the true prevalence of symptomatic recurrent UTI difficult to estimate. Some studies estimate that approximately 44% of women will have recurrence after an initial UTI (2). Regardless, treatment of recurrent UTI results in high costs and significant healthcare burden (5, 6). In post-menopausal women, evidence-based treatment involves the use of topically applied vaginal estrogen and daily prophylactic antibiotics (7). Despite clear efficacy of this regimen (8), there are significant concerns regarding the use of long-term prophylactic antibiotics (9-11).

The pathophysiology of recurrent UTIs is not well understood, but infections are often thought to arise either from repeated ascending bacteria originating outside of the urinary tract, or from reinfection by a persistent reservoir source within the urinary tract (12). One potential reservoir includes the urinary microbiome, which can be quite diverse and contains microbiota considered to be commensals, as well as others that are considered uropathogens (13-16). As such, investigators are keenly interested in understanding how the urinary microbiome may be associated with recurrent UTI pathophysiology. In general, we know that *Lactobacilli* decline in the vaginal niche after menopause, (17) and that vaginal microbiota are highly associated with urinary microbiota (18, 19). Thus, it is not surprising that urinary *Lactobacilli* are also less abundant after menopause (20). This decreased abundance of *Lactobacilli* is thought to play a role in recurrent UTIs. However, it is not clear if non-*Lactobacilli* microbiota may be important as well. Moreover, while *Lactobacilli* in general are relatively resistant to the effects of antibiotics (21), other microbiota within bladder microbial communities (hereafter referred to as the “urobiome”) (22) could be affected by the prophylactic antibiotics used in recurrent UTI management.

We hypothesized that women with recurrent UTIs treated with vaginal estrogen and daily prophylactic antibiotics would have fewer *Lactobacilli* and less microbial diversity compared to age-matched controls without UTIs. Importantly, due to the potentially significant effects of estrogen on bladder microbial communities, we selected control groups who were also using vaginal estrogen for other therapeutic reasons. Thus, we studied three phenotypic groups of post-menopausal women, all concurrently using vaginal estrogen therapy: 1) women with recurrent UTI *on* antibiotic prophylaxis, 2) women with recurrent UTI *not on* antibiotic prophylaxis, and 3) age-matched controls without recurrent UTIs. Our overall objective was to determine whether differences in the urobiome exist between these phenotypic groups. We also aimed to specifically compare the abundance of *Lactobacilli* among groups. We used two complementary techniques to assess microbial communities. The first was expanded quantitative urine culture (EQUC) (23), which allows for detection of live bacteria at the genus and species level. The second was 16S rRNA gene sequencing, a culture-independent method that allows for more thorough characterization of microbiota including fastidious microbes, but often with less resolution at the genus and species level. In order to perform rigorous comparisons, we incorporated Bayesian regression analysis for compositional data, an analytic technique that more accurately models the distribution of microbiome data, incorporates phylogenetic relationships, and allows for the inclusion of other variables to adjust for potential confounders (24).

## RESULTS

### Baseline characteristics

Our study population consisted of 64 women with mean age of 70.5 ± 7.5 years and mean body mass index (BMI) of 29.8 ± 6.3 kg/m^2^. Of these, 17 (27%) were in the recurrent UTI on antibiotic prophylaxis group (rUTI+antibiotics), 24 (38%) were in the recurrent UTI on vaginal estrogen only group (rUTI), while 23 (36%) were in the age-matched control group (controls). There were no significant differences in multiple baseline characteristics (Table 1).

**Table 1:** Baseline characteristics of menopausal women with recurrent urinary tract infections (rUTIs) and healthy controls

|  | rUTI<br>+abx<br>N=17 | rUTI<br>N=24 | Controls<br>N=23 | p-value |
| --- | --- | --- | --- | --- |
| Age (years) | 71.1±7.1 | 71.2±8.7 | 69.3±6.6 | 0.65* |
| BMI (kg/m<sup>2</sup>) | 32.5±8.7 | 29.8±5.7 | 27.8±3.8 | 0.06* |
| Caucasian | 17 (100%) | 23 (95.8%) | 21 (91.3%) | 0.28† |
| African American | 0 (0%) | 1 (4.2%) | 2 (8.7%) | 0.21† |
| Diabetes | 3 (17.6%) | 6 (25.0%) | 1 (4.3%) | 0.12† |
| Daily yogurt consumption | 4 (23.5%) | 9 (37.5%) | 6 (26.1%) | 0.56† |
| Symptoms of fecal incontinence | 2 (11.8%) | 5 (20.8%) | 5 (21.7%) | 0.51† |
| Sexually active | 5 (29.4%) | 9 (37.5%) | 11 (47.8%) | 0.07† |
| Prior incontinence surgery | 4 (25%) | 9 (37.5%) | 9 (39.1%) | 0.45† |
| Prior back surgery | 4 (23.5%) | 3 (12.5%) | 2 (8.7%) | 0.24† |
| History of overactive bladder | 4 (23.5%) | 11 (45.8%) | 6 (26.1%) | 0.74† |

Data are presented as mean±standard deviation, n (%)
Diabetes and BMI were not collinear in our study population (Pearson correlation coefficient 0.19)
rUTI, recurrent urinary tract infection; abx, antibiotics; NA, not applicable; BMI, body mass index
*A priori* pairwise analyses were planned for any significant results that were identified in ANOVA or linear association test. Since none were significant, no pairwise analyses were performed.
\* ANOVA test
† Linear-by-linear association test

### *Lactobacillus* results

Patients were sampled at least 4 weeks after treatment for their most recent infection and in the absence of acute infection. A total of 38/64 revealed growth of at least one microbe in EQUC, with *Lactobacillus* growing in 24/38 (63%) of those exhibiting growth in EQUC. The proportions of urine samples growing *Lactobacillus* in EQUC were not significantly different among groups (29.4% rUTI+antibiotics, 41.7% rUTI, 39.1% controls, χ2 p=0.76). Table 2 shows the individual species that were identified in EQUC when compared to standard urine culture (standard culture did not classify *Lactobacillus* beyond the genus level).

**Table 2:**
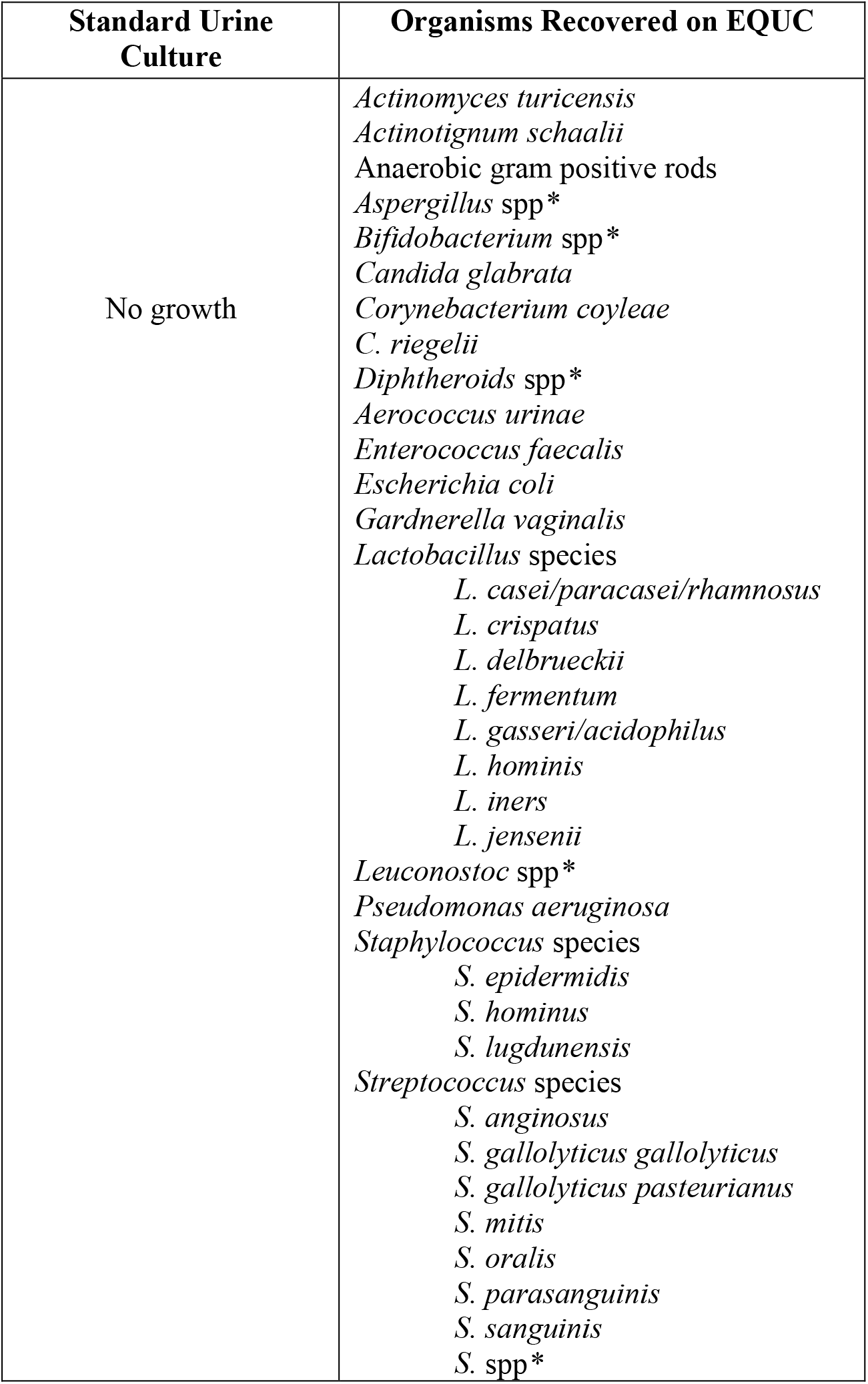
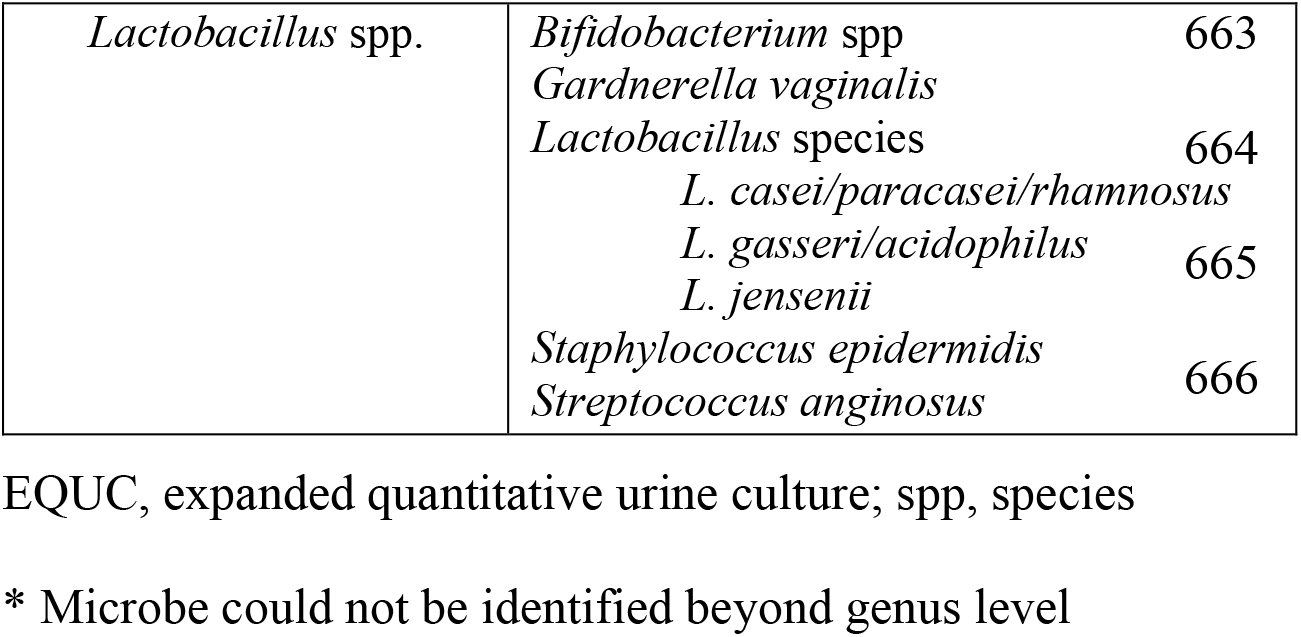
Comparison of organisms recovered using standard urine culture and expanded quantitative urine culture.

| Standard Urine Culture | Organisms Recovered on EQUC |
| --- | --- |
| No growth | *Actinomyces turicensis*<br>*Actinotignum schaalii*<br>Anaerobic gram positive rods<br>*Aspergillus* spp*<br>*Bifidobacterium* spp*<br>*Candida glabrata*<br>*Corynebacterium coyleae*<br>*C. riegelii*<br>*Diphtheroids* spp*<br>*Aerococcus urinae*<br>*Enterococcus faecalis*<br>*Escherichia coli*<br>*Gardnerella vaginalis*<br>*Lactobacillus* species<br>*L. casei/paracasei/rhamnosus*<br>*L. crispatus*<br>*L. delbrueckii*<br>*L. fermentum*<br>*L. gasseri/acidophilus*<br>*L. hominis*<br>*L. iners*<br>*L. jensenii*<br>*Leuconostoc* spp*<br>*Pseudomonas aeruginosa*<br>*Staphylococcus* species<br>*S. epidermidis*<br>*S. hominus*<br>*S. lugdunensis*<br>*Streptococcus* species<br>*S. anginosus*<br>*S. gallolyticus gallolyticus*<br>*S. gallolyticus pasteurianus*<br>*S. mitis*<br>*S. oralis*<br>*S. parasanguinis*<br>*S. sanguinis*<br>*S. spp\** |

|  |  |  |
| --- | --- | --- |
| *Lactobacillus* spp. | *Bifidobacterium* spp | 663 |
|  | *Gardnerella vaginalis* |  |
|  | *Lactobacillus* species | 664 |
|  | *L. casei/paracasei/rhamnosus* |  |
|  | *L. gasseri/acidophilus* | 665 |
|  | *L. jensenii* |  |
|  | *Staphylococcus epidermidis* | 666 |
|  | *Streptococcus anginosus* |  |

EQUUC, expanded quantitative urine culture; spp, species
\* Microbe could not be identified beyond genus level

Next, we performed logistic regression analyses to control for potential confounders and to assess for factors independently associated with the presence of *Lactobacillus* growth in EQUC. Though history of diabetes was associated with increased odds of growing *Lactobacillus* in EQUC (OR 5.56, 95% CI 1.14-27.02), when controlling for this variable as well as BMI and sexual activity, there were still no differences in growth of *Lactobacillus* in EQUC between study groups.

We further analyzed the most prevalent *Lactobacillus* species recovered (*L. gasseri/acidophilus, L. casei/paracasei*, and *L. crispatus*) in EQUC. We calculated the proportions of samples with growth of these species and compared proportions between study groups. There were trends toward fewer *L. gasseri* and *L. cripatus* in the rUTI+antibiotics group, but these did not reach statistical significance (Figure 1a).

**Fig 1a:**
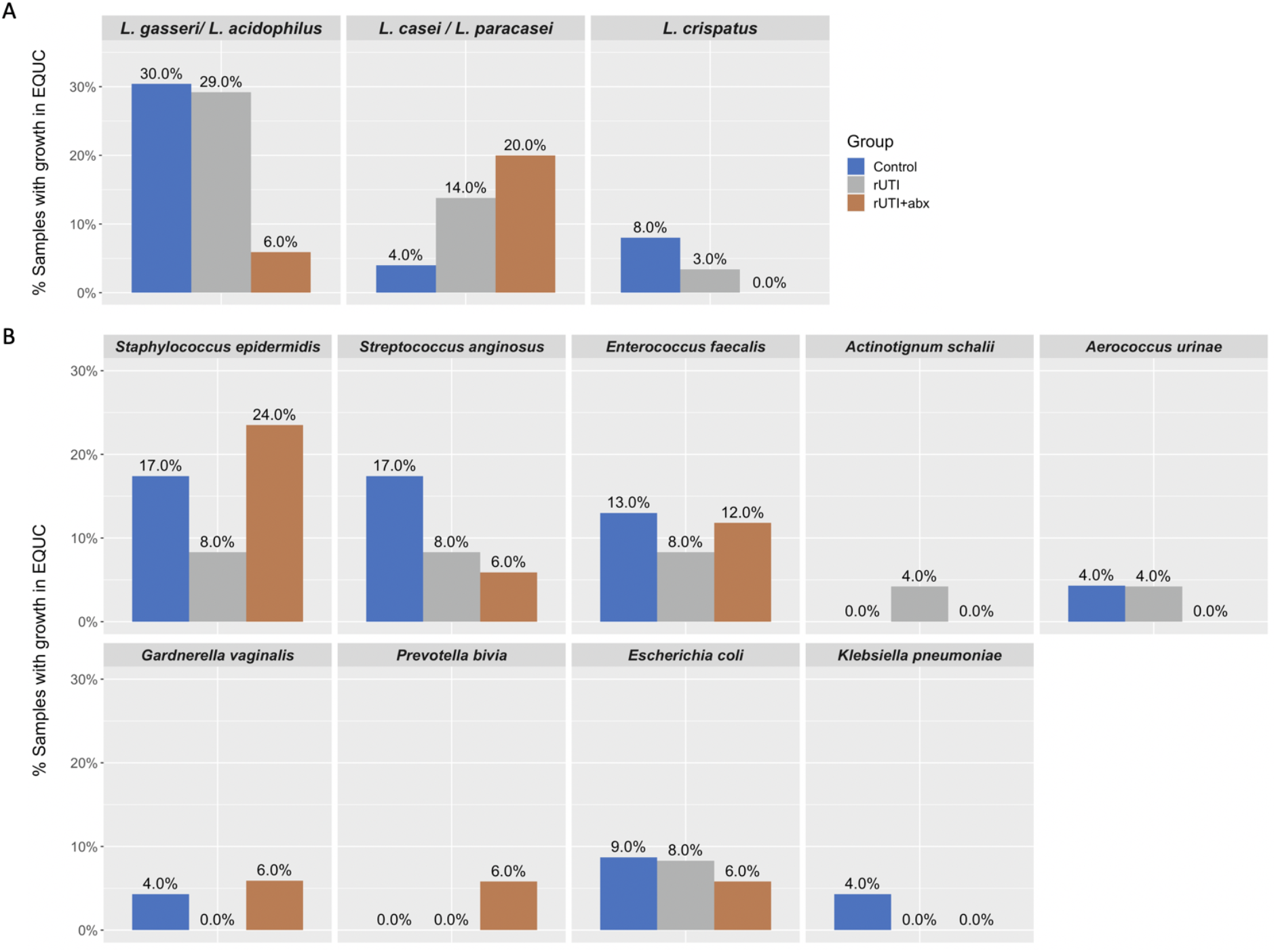
Comparison of the most common *Lactobacillus* organisms recovered in EQUC among groups. Of 64 total samples, 38 (59%) demonstrated growth of one or more organisms in expanded quantitative urine culture (EQUC). The majority of these organisms were *Lactobacilli*. The three most commonly isolated *Lactobacillus* species were L. *gassed*/*L. acidophilus* (isolated in n=15 samples), *L. casei*/*L. parpcasei* (n=6 samples), and *L. crispatus* (n=3 samples). Proportions of urine samples from each phenotypic group with the three most common *Lactobacillus* species in EQUC are depicted in the figure. For example, in women with recurrent UTI taking antibiotics, 6% contained L. *gasseri*/*L. acidophilus* in their EQUC samples compared to 29-30% in the other two phenotypic groups. These differences in proportions were not statistically significant (p=0.14, p=0.39, p=0.43, respectively). **Figure 1b: Comparison of other organisms recovered on EQUC among groups**. A total of 38 urine samples grew one or more organisms in expanded quantitative urine culture (EQUC), with the most common non-*Lactobacillus* organisms identified here. Proportions of urine samples from each phenotypic group where the microbe was recovered in EQUC are depicted in the figure. There were no statistically significant differences.

With regard to microbiota identified through 16S rRNA sequencing, recovered taxa were highly variable with significant variability among groups (Figure 2). The family *Lactobacillaceae* were recovered in 97% of samples. Due to the sequencing strategy that utilized a single region of the 16S rRNA gene, we were not able to taxonomically classify *Lactobacillaceae* at higher resolutions to determine individual genera and species. We further categorized samples as having “low”, “moderate”, and “high” relative abundances of *Lactobacillaceae* using thresholds of <20% to denote “low” and >80% to denote “high” abundance. In other words, if <20% of the total microbiota recovered from a sample were from the family *Lactobacillaceae* compared to other families, this was considered “low” abundance. Based on this threshold, there was moderate or high *Lactobacillaceae* abundance in 6/17 (35%) of samples in the rUTI+antibiotics group, 9/24 (37.5%) of samples in the rUTI group, and 13/23 (56.5%) of samples in the control group, with no statistically significant difference among groups (χ2 p=0.3) (Figure 3). None of the samples in the rUTI+antibiotics group achieved “high” abundance compared to 4/24 (16.6%) and 5/23 (21.7%) in the rUTI and control groups, respectively.

**Figure 2:**
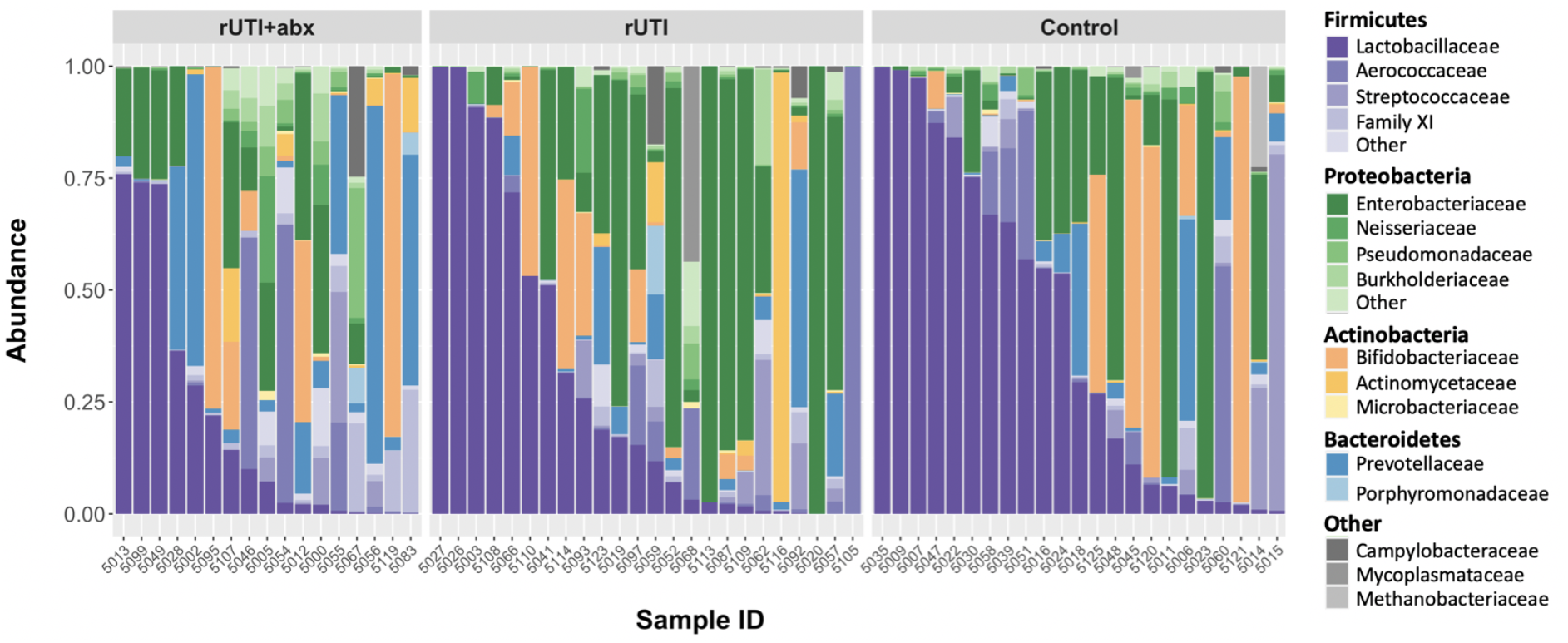
Stacked bar plot depicting relative abundance of all microbiota per sample. Each vertical bar depicts the relative abundance of adjusted sequence variants (ASVs) and associated taxa that were recovered per sample. For each color group, the darkest shade represents the most common family identified with a phylum, with lighter shades representing other families within the same phylum. Results obtained from V4 amplicon sequencing of the 16S rRNA gene, processed with DADA2 and mapped to the SILVA reference database.

**Figure 3:**
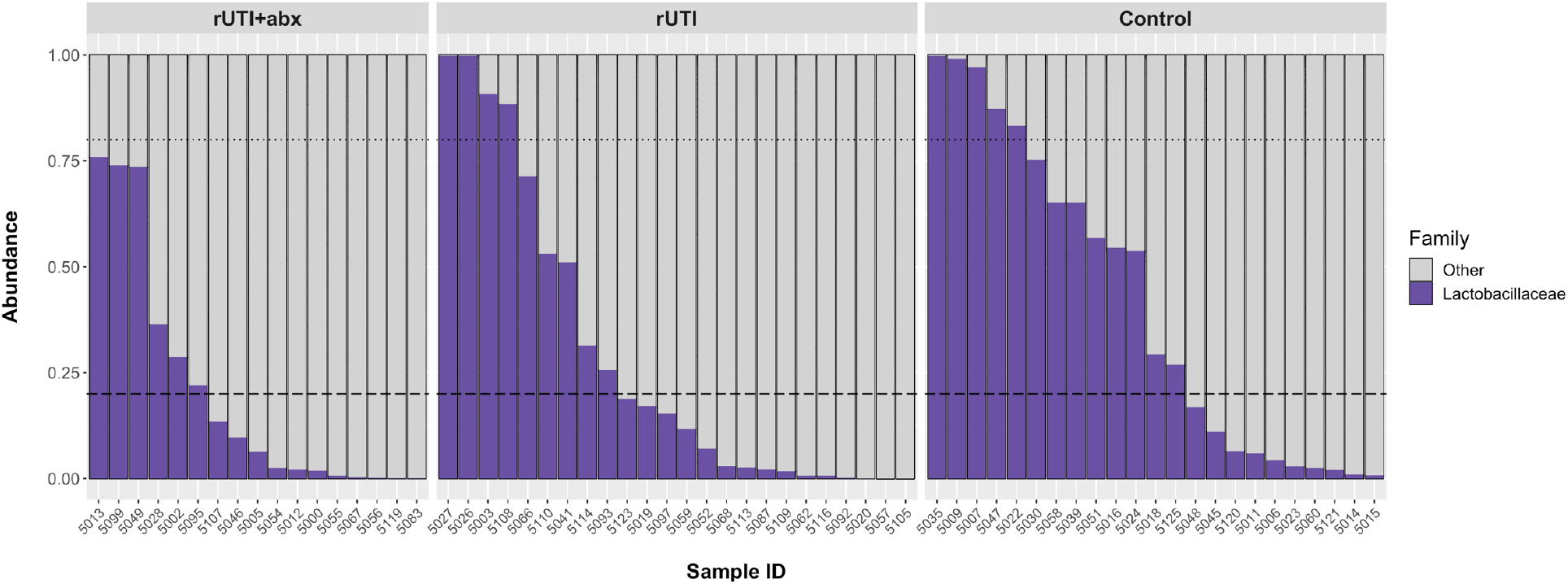
Stacked bar plot depicting relative abundance of *Lactobacillaceae* per sample. Stacked bar plot where each vertical bar depicts the relative abundance of the family *Lactobacillaceae* recovered per sample (colored purple), with all other taxa colored gray. The dashed line indicates the 20% threshold; samples with bars below this line are considered to have “low” relative abundance of *Lactobacillaceae*. The dotted line indicates the 80% threshold; samples above this bar are considered to have “high” relative abundance of *Lactobacillaceae*.

### Other microbial results

There were no significant differences among groups in the presence or median numbers of microbial species that grew in EQUC, with 2 [IQR 1,2.5] in the rUTI+antibiotics group, 1 [IQR 1,2] in the rUTI group, and 1 [IQR 1,3] in the control group (Kruskal-Wallis test p=0.91). Thirty-eight women (59.4%) had at least one microbial organism recovered on EQUC in the setting of a “no growth” standard urine culture (Table 2).

We compared the proportions of women whose urine samples grew known or potential uropathogens in EQUC (*Staphylococcus epidermidis, Streptococcus anginosus, Enterococcus faecalis, Actinotignum schalii, Aerococcus urinae, Gardnerella vaginalis, Prevotella bivia, Escherichia coli and Klebsiella pneumoniae*). There were no statistically significant differences among groups when comparing the proportions with growth of these bacteria (Figure 1b).

We next assessed 16S rRNA sequencing data to further compare the urobiome microbial communities. Non-metric multidimensional scaling (NMDS) techniques were applied to cluster samples based on Unifrac distance without significant differences (Supplemental Figure 1). We then performed permutational multivariate analysis of variance (PERMANOVA) comparisons of Unifrac distances while controlling for several covariates. In this PERMANOVA model, we identified significant differences in Unifrac distances, or in other words, differences in biological communities, among the three groups (p=0.045). Of the covariates, sexual activity was significantly associated with increased Unifrac distance (p=0.04), while age and BMI approached statistical significance (p=0.05 and p=0.06, respectively, Supplemental Table 1).

Pairwise comparisons were performed to further understand which specific groups were driving the overall differences in the urobiome. In these pairwise comparisons, PERMANOVA models were again constructed to compare Unifrac distances between groups while adjusting for the same covariates as above. There were significant differences in microbial communities between women with rUTI + antibiotics compared to controls (p=0.038) where again, sexual activity was the only covariate significantly associated with the outcome (p=0.005). Other pairwise comparisons showed non-significant differences in PERMANOVA analyses (rUTI+antibiotics vs. rUTI, p=0.086; rUTI vs. controls p=0.12).

PERMANOVA analyses can identify differences in overall compositions, but have limited ability to detect the group of taxa that are responsible for those differences. Also, because of the high cross-sample variability of bacterial taxa found within the urinary microbiome, traditional models such as PERMANOVA have limited power to separate signal from noise. Thus, we also performed Bayesian graphical compositional regression (BGCR) modeling, which is a technique that assesses all of the branches of the phylogenetic tree to see if there are statistically significant differences between groups at each split of the phylogenetic tree after adjusting for covariate effects. Though we are limited to only pairwise comparisons, we are able to drill down to specific taxa that differ between phenotypic groups, while incorporating covariates. BGCR calculates the posterior probability for whether the overall microbial compositions differ, which is referred to as the posterior joint alternative probability (PJAP). A value of PJAP exceeding 0.5 provides evidence for the presence of cross-group difference not due to multiple testing, with larger PJAPs (up to 1) indicating stronger evidence. BGCR also calculates a posterior probability for whether the groups differ at each split of the phylogenetic tree, which we refer to as the posterior marginal alternative probability (PMAP) at that split. We explored splits where the PMAP was >60%.

The BGCR analysis is generally consistent with the results of PERMANOVA. When comparing women with rUTI +antibiotics to controls, where significant differences were noted in PERMANOVA analyses, there is again a high overall probability of differences in microbial composition (PJAP = 99.9%) when analyzed with BGCR. There are five branches of the phylogenetic tree where the probability of a difference is quite high (Fig 4, Supplemental Figure 2). Three of these branches are downstream branches of each other and identify high probabilities of differences in the genus *Prevotella*, family *Prevotellaceae*, and order Bacteroidales (PMAP = 99.9%, 82.5%, and 80.5% respectively) between groups. Additional distinguishing features include high probabilities of differences in the order Clostridiales, family *Ruminococcaceae* (PMAP = 84.8%), and in different Actinobacteria (PMAP = 98.3%), which are summarized in Supplemental Figure 2.

Slightly different than in the PERMANOVA analyses, when comparing women with rUTI + antibiotics to those with rUTI not using antibiotics, the overall probability of differences in microbial composition is relatively high with BGCR (PJAP = 82.9%). Of the various splits in the phylogenetic tree, the highest probability of differences occurred within the order Clostridiales, family XI (PMAP = 66.7%, Fig 5), where different Clostridial genera were noted identified in women with rUTI + antibiotics compared to those with rUTI not using antibiotics.

**Figure 4:**
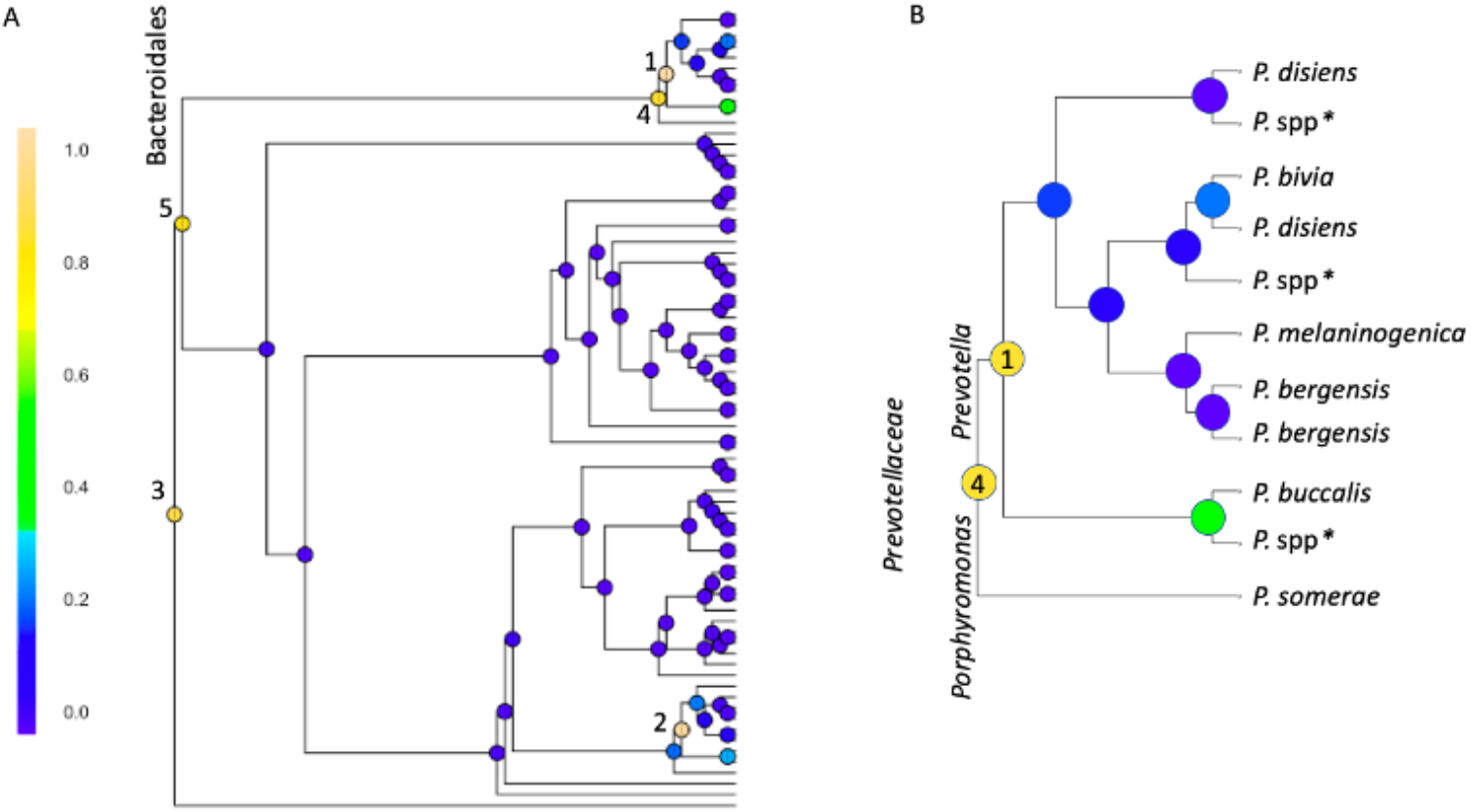
Bayesian GCR analysis comparing urine microbiota in women with recurrent UTIs taking daily antibiotics compared to age-matched controls (both using vaginal estrogen). Results of BCGR analysis comparing the two groups while incorporating multiple clinical and technical covariates. The posterior marginal alternative probability (PMAP) was calculated at each node and shaded based on result. The highest probability of a difference between groups is at node #1 (PMAP = 99.96%), which denotes differences in *Prevotella* species recovered among those with rUIJ+abx compared to controls. This node is a downstream branch in the taxonomic tree from two other nodes that also appear to have a high probability of differences between groups. These are identified as #4 (PMAP = 82.5%), which denotes differences in *Prevotellaceae* and #5 (PMAP = 80.5%), denoting differences in the order Bacteriodales. Node #2 (PMAP = 98.27%) denotes differences in Actinobacteria; this section of the tree is further exploded in Supplemental Figure 2. Node #3 (PMAP = 84.81%) denotes differences in Clostridiales, family *Ruminococcacege*. Fig 4A shows the entire taxonomic tree and BCGR results; Figure 4B is an exploded section of this tree showing genus and species information.

**Figure 5:**
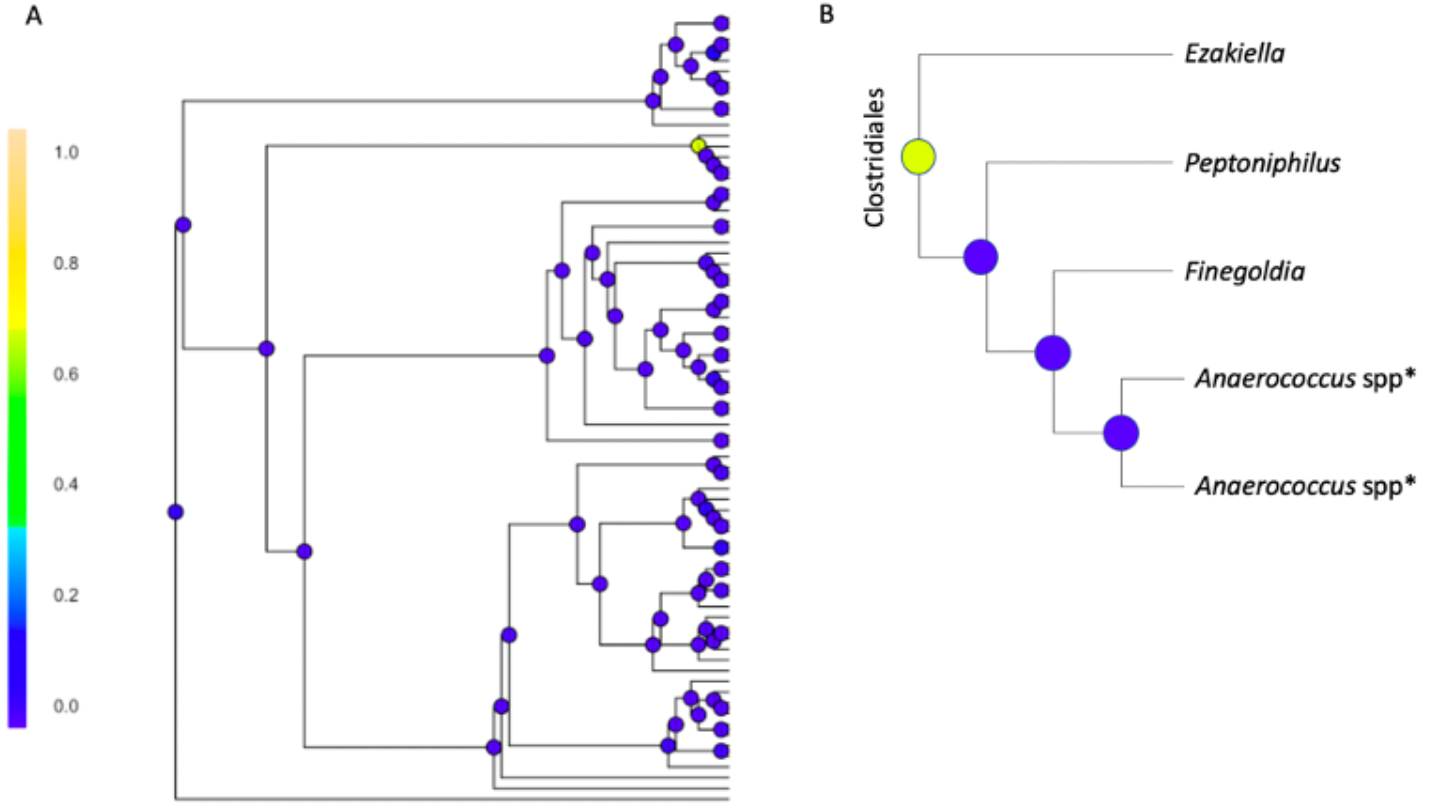
Bayesian GCR analysis comparing urine microbiota in women with recurrent UTIs taking daily antibiotics compared to women with recurrent UTIs not on antibiotics (both using vaginal estrogen), (a) Posterior marginal alternative probability (PMAP) was calculated at each node with one area resulting in PMAP > 60%. This corresponds to the node shaded in yellow-green where PMAP = 66.73%, indicating the probability of a true difference in taxa between groups at the location specified, (b) Exploded view of the taxa where the difference was identified, specifically showing that there are differences between the genus *Ezakiella*, and the other genera identified in the figure.

When comparing women with rUTI who are not using antibiotics to matched controls, there was only a modest probability of an overall difference (PJAP = 63.8%) with no individual branch splits showing probabilities of differences (all PMAP < 55%). These results echo those found with PERMANOVA where there was a lack of significant differences between groups.

Finally, we assessed the concordance of microbiota identified from the same sample through EQUC compared to 16S rRNA sequencing. We assessed concordance at the family level since 16S rRNA did not always identify taxa at the genus or species level (Fig 6). As suspected, though some microbes were identified with both techniques, many more were only identified through 16S rRNA sequencing. The families *Lactobacillaceae* and *Streptococcaceae* had the highest concordance, and yet they were only approximately 50% concordant. Notably, there were a few families, including *Enterococcaceae*,and *Corynebacteriaceae*, where organisms only grew in culture and were not identified by sequencing.

**Figure 6:**
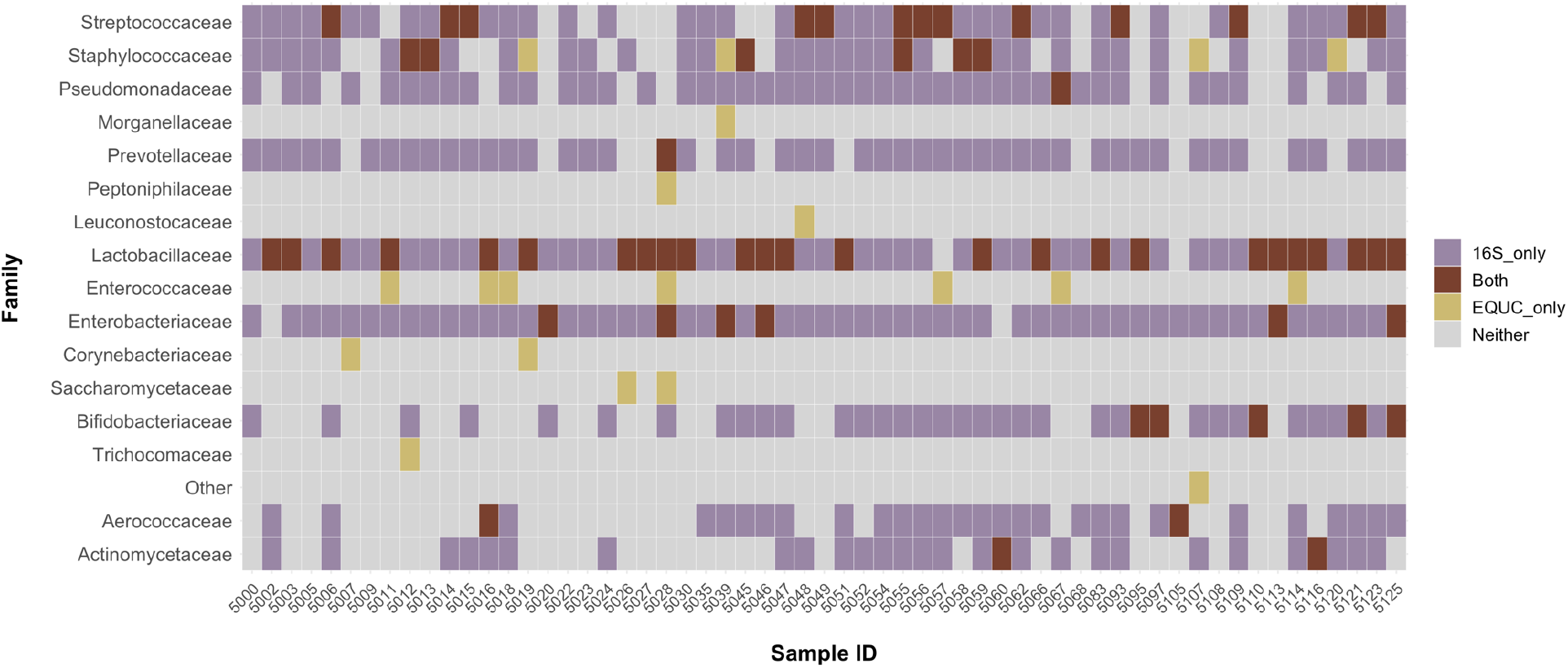
Correlation of expanded quantitative urine culture (EQUC) and 16S rRNA sequencing results. This figure depicts the various families of organisms that were recovered on expanded quantitative urine culture (EQUC) and 16S rRNA sequencing within each sample. Organisms are depicted in the rows and samples are depicted in columns. The colored bars denote whether each organism was recovered on 16S rRNA sequencing only, both methods, EQUC only, or neither method.

## DISCUSSION

In this cross-sectional analysis of the urobiome in menopausal women using vaginal estrogen, we did not identify significant differences in overall abundance of *Lactobacilli* among those with recurrent UTIs and matched controls. However, there were trends towards different *Lactobacillus* species among groups. In women with recurrent UTIs taking daily antibiotics, multiple analytic techniques (e.g. PERMANOVA and BGCR) identified differences in microbial compositions when compared to controls without recurrent UTIs. A BGCR analysis shows that these differences are mainly attributed to anaerobic bacteria including differences in *Prevotellaceae*, Clostridiales, as well as the class Actinobacteria. BGCR analysis, which is more precise than traditional analyses such as PERMANOVA, also uniquely identified a high probability of differences in Clostridial genera between women with recurrent UTIs taking daily antibiotics compared to those not taking daily antibiotics.

There are a number of strengths inherent to our study. Due to purposeful sampling, we were able to standardize the timing of sample acquisition (>4 weeks after acute UTI, and >6 weeks after initiating estrogen and/or prophylactic antibiotics) and collect information on variables that can significantly confound microbial data. Importantly, we standardized the use of vaginal estrogen, which is a large potential confounder for the urinary microbiome, and also excluded medications that could potentially influence the urobiome. All women underwent catheterized sampling, which allows confidence that we are characterizing bladder microbiota, as described in prior studies (16). We used complementary techniques of expanded quantitative urine culture (EQUC) and 16S rRNA gene sequencing to characterize the urobiome. The 16S rRNA sequencing data allow for broader identification of microbiota and more precise conclusions regarding bacterial phylogenetic relationships among the groups, while the EQUC data provide important information about *viable* urinary microbiota, as well as species-level information. We also focused on clinically relevant recurrent UTI groups (the first on antibiotic prophylaxis, the second not using antibiotics), where data characterizing the urobiome are currently sparse.

The primary limitations of this study are inherent to the small sample size and cross-sectional design. Despite having a large clinical population of women with recurrent UTIs, we had strict entry criteria including verification of culture-proven UTIs and exclusion of women taking UTI prevention supplements like methenamine, cranberry tablets, probiotics and D-mannose. These factors ultimately restricted the number of potential candidates for this study. There are multiple findings, for example those pertaining to *Lactobacillus* abundance and differences in *Lactobacillus* species, where we can see trends towards differences in groups that do not reach statistical significance, possibly due to the small sample size. Despite the small sample size, we still observed some significant differences among phenotypic groups and thus we believe this analysis still provides useful information. However, many of our conclusions relate to the presence of anaerobic bacteria in the urine of women with recurrent UTIs taking daily prophylactic antibiotics. Due to the cross-sectional sampling we are unable to determine if these differences are due to selective pressure from prolonged antibiotics, or if these anaerobes are implicated in the development of more severe recurrent UTIs that require prophylactic antibiotics. Notably, we did not standardize the antimicrobial agent used for prophylaxis, as that would have further hindered enrollment. Antibiotics were chosen based on clinical indicators including allergies, resistance patterns, and interactions with other medications. Though the majority of patients were taking trimethoprim as a daily prophylactic antibiotic, some were also taking nitrofurantoin or cephalexin. Thus, the different mechanisms of antimicrobial activity could also influence our findings in women taking daily antibiotics.

One microbe of interest in this study was *Lactobacillus* (20, 25). In general, *Lactobacilli* are relatively resistant to the effects of antibiotics (21), but tend to decline in the absence of estrogen (20). As such, it is not surprising that microbes from the family *Lactobacillaceae* were found in 97% of our samples, since all women were using vaginal estrogen therapy. Though not statistically significant, the species-level results pertaining to *Lactobacilli* could be biologically relevant. Prior data show that *L. gasseri* comprise one *Lactobacillus* species that is likely to secrete bacteriocins (26), or bacterial toxins that antagonize other bacteria. In our study, women with recurrent UTIs taking daily antibiotics had a disproportionately low amount of *L. gasseri* compared to the other phenotypic groups. These findings support the theory that the presence of specific *Lactobacillus* species, rather than general *Lactobacillus* abundance, may be important in the pathophysiology of UTIs. Similarly, our data suggest that prophylactic antibiotics are associated with shifts in which *Lactobacillus* species are present in the urobiome.

Our data contribute to the growing body of literature regarding the urobiome and urologic disorders. Furthermore, we utilized BGCR, a published analytic technique (24), designed to identify individual taxa contributing to phenotypic differences. Using this technique, we are able to hone in on specific taxa that can be explored in future mechanistic studies. We also found associations between microbial composition and several covariates including sexual activity, age, BMI and history of diabetes. Many of these covariates are also associated with recurrent UTIs and could serve as significant confounders of microbial results, if not incorporated into analyses. In addition to presenting new ways to analyze microbial data, our findings confirm the importance of adjusting for important clinical covariates within studies of the urobiome. Finally, we also present a novel analysis comparing EQUC and 16S rRNA data from the same sample, collected at the same time. This concordance analysis highlights the limitations of culture-based methods to identify urinary bacteria. However, there were a few bacterial families, including *Entercococcaceae* which are relevant for UTIs, that were only identified using EQUC and not through sequencing. In exploring the microbes that were not recovered with sequencing, the majority were Gram positive bacteria or yeast/fungi. Notably, these organisms have a thicker cell wall that is harder to lyse, and sequence-based techniques for identification of microbes are known to have biases towards Gram negative bacteria. As such, special procedures such as additional mechanical or enzymatic methods of cell lysis, should be employed if investigators are interested in thoroughly understanding the Gram-positive component of a microbial community.

Other groups have reported evidence for synergistic interactions amongst urinary and other microbes that enhance bacterial colonization and/or persistence (12, 27). Our findings overall support the idea that it may not be one particular microbe that puts a woman at risk for recurrent UTIs, but perhaps a shift in the balance of *Lactobacilli*, or particular species of *Lactobacilli* compared to anaerobic bacteria. Our findings suggest that future studies of recurrent UTI should consider complex interactions among bacteria to identify the components of the urobiome most associated with bladder health, and should also include a longitudinal component in order to explore changes in the urobiome over time.

## MATERIALS AND METHODS

### Clinical Procedures and Sample Acquisition

After approval from the Institutional Review Board at Duke University Medical Center (IRB # Pro00083917), we performed a translational study of women from the Female Pelvic Medicine and Reconstructive Surgery (FPMRS) and Urology clinics at Duke University Health System from December 2017 to March 2019. This cross-sectional analysis was nested within a larger longitudinal study of the urinary microbiome in women with recurrent UTI. Menopausal women using vaginal estrogen from three groups were included: 1) Recurrent UTI *on* antibiotic prophylaxis (rUTI+antibiotics); 2) Recurrent UTI *not on* antibiotic prophylaxis (rUTI); and 3) Age-matched controls without recurrent UTIs (controls). Women aged 55 years and older were included to ensure menopausal status. All participants were required to have been on vaginal estrogen therapy (cream, tablet or ring) at least once weekly for a minimum of 6 weeks prior to enrollment. Within the previous 24 months, women in the two recurrent UTI groups were required to have met diagnostic criteria of two or more culture-proven symptomatic UTIs within a 6-month period, or three culture-proven UTIs within a 12-month period. Exclusion criteria included currently using UTI prevention supplements (methenamine, probiotics, D-mannose, cranberry tablets) and not willing to discontinue 6 weeks prior to urine sampling, instrumentation of urinary tract in previous month, neurogenic bladder or any other need for chronic intermittent catheterization, history of prior pelvic radiation, active malignancy, breast cancer within previous 5 years or taking any anti-estrogen medication (e.g. aromatase-inhibitors or selective estrogen receptor modulators), history of neurologic condition that may affect urinary function (e.g. stroke, multiple sclerosis, spinal cord injury, Parkinson’s disease), known renal insufficiency with creatinine > 1.3, immunosuppression or chronic steroid use, intravaginal pessary use, using only post-coital antibiotic prophylaxis (excluded from all groups), treatment with antibiotics for any active infection in the previous month. For the rUTI +antibiotics group, acceptable antibiotics included trimethoprim, nitrofurantoin, or cephalexin used daily for at least 6 weeks prior to urine sampling.

For this analysis, we used a convenience sample of the first 64 women enrolled in a larger longitudinal study. This pilot study was designed to explore hypotheses, feasibility, and lay the groundwork for larger, hypothesis testing studies. At the first sampling visit, informed consent was obtained, and demographic and medical history were collected. Participants completed validated questionnaires to assess for pelvic floor symptoms including overactive bladder, urinary incontinence, and fecal incontinence (Pelvic Floor Distress Inventory, PFDI-20) (28). A non-validated questionnaire was also administered to assess for special diets, types of foods and beverages ingested on a daily basis, and sexual activity. A catheterized transurethral urine sample was obtained by clinical personnel. Chemical urinalysis was performed; if there was evidence of active UTI (> 1+ blood, > 1+ leukocytes or + nitrites) in the presence of UTI symptoms, the sample was sent for standard urine culture for diagnostic purposes only. The participant was treated with antibiotics as indicated based on standard urine culture and asked to return in 4 weeks for a repeat sampling visit. If the participant had another symptomatic UTI at the repeat sampling visit, they were given a third and final sampling visit opportunity. Those with symptomatic UTI at three sampling visits in a row were excluded. Other reasons to delay urine sampling for 6 weeks included: need to start vaginal estrogen or prophylactic antibiotic and discontinuation of UTI prevention supplements/medications including cranberry, probiotic, D-mannose, or methenamine.

### Sample Processing

If there was no evidence of UTI either by symptoms or chemical urinalysis, then the urine sample was divided into two portions. The first portion was placed within two 5 mL BD Vacutainers® using sterile technique. Both culture vacutainers were transported on ice via next available courier to the Clinical Microbiology Laboratory at Duke University and processed within 24h of collection. One culture vacutainer was sent for standard urine culture, while the other was sent for expanded quantitative urine culture (EQUC). The protocol for EQUC has been published elsewhere (29). In brief, EQUC involves a series of modifications to the standard urine culture, such as additional culture media, longer incubation time, larger volume of urine, and varied atmospheric conditions (including anaerobic). These modifications allow for growth of fastidious or anaerobic organisms, as well as the detection of organisms that are present in small numbers. Bacterial species identification was performed by the clinical microbiology laboratory via matrix-assisted laser desorption/ionization – time of flight (MALDI-TOF) mass spectrometry.

The second portion (reserved for 16S rRNA gene sequencing) of urine was sterilely placed within 50 ml 10% Assay Assure tubes (Thermo Fisher Scientific; Waltham, MA) and transported on ice via next available courier to our shared laboratory space on the Duke University campus, where samples were refrigerated at 4°C for no more than 96 hours prior to initial processing. For initial processing, the Assay Assure tubes underwent a series of centrifugations and washing with phosphate-buffered saline. The final dry urine pellets were stored at -80°C until all samples were collected.

### DNA Isolation and Sequencing

Once all urine pellets were collected, DNA extraction was performed using the cultured cell protocol of the Qiagen DNeasy Blood and Tissue Kit with the addition of the optional RNase step (Valencia, CA, product code # 69506) according to manufacturer instructions (30). DNA was eluted into 50 μL of buffer AE and stored at -20°C. Next, we prepared a series of samples that would serve as positive and negative controls for sequencing. For negative controls, we prepared one tube of 50 μL microbial-free water. For the positive controls, we prepared a series of 8 serial dilutions of a known mock microbial community consisting of 8 gram positive and negative bacteria and 2 yeast (Zymo Research, catalog # D6300), as previously described (31). The dilutions were prepared in a 1:2 ratio with microbial free water. All positive and negative controls were subjected to the same DNA extraction steps using the same reagents as the study samples, and elution with 50 μL of buffer AE. DNA quality was assessed using Nanodrop spectroscopy to calculate the 260/280 ratio (Thermo Fisher Scientific; Waltham, MA), and DNA was quantified in ng/μL with Qubit fluorometry (Thermo Fisher Scientific; Waltham, MA). The samples were submitted to the Duke Microbiome Shared Resource for library preparation and 16S rRNA gene sequencing.

Briefly, a 16S rRNA gene amplicon library was generated via PCR amplification of the V4 hypervariable region of the 16S rRNA gene using the forward primer 515 and reverse primer 806 following the Earth Microbiome Project protocol (http://www.earthmicrobiome.org/). Equimolar PCR products from all samples were pooled prior to sequencing. Sequencing was performed by the Duke Sequencing and Genomic Technologies shared resource on an Illumina MiSeq instrument configured for 250 base-pair paired-end sequencing runs as published elsewhere (29).

### Bioinformatics & Statistical Analysis of Sequencing Data

Raw sequences were processed into amplicon sequence variants (ASVs) with DADA2 (v 1.14.0) (32) using default parameters except for the trim and trunc parameters which were 20 (trim), 190 (trunc reverse reads), and 230 (trunc forward reads). ASVs were mapped to the SILVA reference database (v 132) for taxonomic identification using the RDP classifier implemented in DADA2. Decontam (v 1.2.1) was used to identify potential contaminant ASVs using the frequency method and a threshold of 0.5 (33), which was chosen after evaluating the contaminant removal of the mock microbial dilution series (31). Data were further processed and visualized in R using phyloseq (v. 1.26.1) (34).

Prior to statistical analyses, an additional 2-step filtering process was used to further remove noise from the data; amplicon sequence variants (ASVs) below a relative abundance threshold of 0.001 were removed, and ASVs of < 1% prevalence in the entire dataset were also removed. We explored samples whose DNA had a 260/280 ratio of < 1.6 on Nanodrop spectrophotometry (n=9) to ensure that DNA sample quality was not affecting downstream results. Ultimately there were no differences in overall findings, and no major differences in microbial findings among samples with 260/280 ratio of < 1.6 compared to those with higher ratios; thus all samples were retained. Data were then re-normalized such that the relative abundance of the ASVs in each sample sum to 1. Recovered sequences were plotted to compare taxonomic structure and stacked bar plots were reviewed. For the family *Lactobacillaceae*, we categorized samples based on abundance. Though thresholds for “low” and “high” abundance have not been formally specified, we used thresholds that are consistent with other reports in the literature referring to low and high *Lactobacillus* abundance in urine (20, 25).

We used non-metric multidimensional scaling (NMDS) to cluster microbial communities between groups based on Unifrac distance. Unifrac distances were compared among the three patient groups using permutational multivariate analysis of variance (PERMANOVA) while incorporating clinical covariates (age, BMI, diabetes, daily yogurt ingestion, overactive bladder, fecal incontinence, sexual activity) and technical covariates (260/280 ratio from Nanodrop spectrophotometry, logarithmically transformed DNA concentration, urine pH, urine specific gravity, and duration of time from sample collection to final processing). These covariates were chosen based on clinically relevant variables as well as those that were statistically significant in univariate analysis. Further pairwise comparisons were performed with PERMANOVA with the same covariates, followed by Bayesian graphical compositional regression analysis (BGCR) (24), also with the same covariates. BGCR returns the posterior joint alternative probability (PJAP), which summarizes the overall test result of the null hypothesis that the microbiome composition of the two comparison groups are the same. It carries out a comparison of the subcomposition of taxa at each split of the phylogenetic tree to examine the cross-group difference and combines the information across the entire tree through a graphical model thereby enhancing its power to identify small differences through borrowing of strength over the tree. It incorporates a multiple testing control strategy by specifying the graphical model such that the prior alternative probability, or the probability for the presence of any cross-group difference in at least one split of the phylogenetic tree, does not exceed 0.5 regardless of the size of the tree or the total number of splits. Therefore, a value of PJAP exceeding 0.5 provides empirical evidence for the presence of cross-group difference after multiple testing adjustment, with larger PJAPs (up to 1) indicating stronger evidence. BGCR also returns the posterior marginal alternative probability (PMAP) at each split of the phylogenetic tree, that quantifies the empirical evidence for the presence of cross-group difference in the subcomposition at that split. We qualitatively explored the descendant ASVs at any split with PMAP >0.6 (i.e. a 60% or greater probability that there are differences in the taxa at that phylogenetic branch between groups).

Finally, we plotted the proportion of samples having growth of a variety of microbes by family on EQUC and correlated this with the proportion of samples that were sequence positive for the same families on 16S rRNA gene sequencing.

### Other statistical analyses

Demographic and baseline characteristics were compared between the three groups. Continuous variables were analyzed using one-way analysis of variance (ANOVA) and t-tests, as appropriate. Categorical variables were analyzed using chi-square, Fisher’s exact and Linear-by-Linear association tests, as appropriate. The median number of microbial species recovered from EQUC was compared among the three groups using the Kruskal-Wallis test. Multivariable logistic regression was performed to adjust for potential confounders when assessing EQUC results. A first logistic regression model was created for variable selection and included theoretical risk factors for dysbiosis of urinary microbial flora (Supplemental Table 2). The final regression model included BMI, sexually active status and history of diabetes mellitus as covariates, as these were the only covariates that had a p-value of <0.1 in either the full regression model or bivariate analysis. Data were analyzed using IBM SPSS Statistics version 24.0 (SPSS Inc., Armonk, NY) and R packages phyloseq (version 1.32.0) and BGCR (version 0.1.0).

## ACKNOWLEDGEMENTS

The authors would like to thank the following research coordinators from the Duke Division of Urogynecology and Reconstructive Pelvic Surgery for their valuable contributions: Akira Hayes, Acacia Harris, Shantae McLean, Robin Gilliam, Claire McLaughlin, and Yasmeen Bruton. The authors would also like to thank Carole Grenier, research analyst from the Murphy Laboratory at Duke University, for her expert contributions to sample preparation for DNA extraction. Library preparation and DNA sequencing were performed in collaboration with the Duke Microbiome Center Core Facility. Dr. Kevin C. Hazen from the Duke Clinical Microbiology laboratory provided insightful contributions and staff assistance with expanded quantitative urine culture. Finally, Erin Dahl from Oregon Health & Science University assisted with creating figures for this manuscript.

## PUBLIC DATA SHARING

All sequences are available for download in the Sequence Read Archive under Accession Number PRJNA685466 (http://www.ncbi.nlm.nih.gov/bioproject/685466)

